# Generative Adversarial Networks (GAN) for the simulation of central-place foraging trajectories

**DOI:** 10.1101/2021.09.27.461940

**Authors:** Amédée Roy, Sophie Lanco Bertrand, Ronan Fablet

**Affiliations:** Institut de Recherche pour le Développement (IRD), MARBEC (Univ. Montpellier, Ifremer, CNRS, IRD), Avenue Jean Monnet, 34200, Sète, France; IMT Atlantique, UMR CNRS Lab-STICC Brest, France

**Keywords:** Animal telemetry, Deep Learning, Hidden Markov Model, Movement model, Seabird

## Abstract

1. Miniature electronic device such as GPS have enabled ecologists to document relatively large amount of animal trajectories. Modeling such trajectories may attempt (1) to explain mechanisms underlying observed behaviors and (2) to elucidate ecological processes at the population scale by simulating multiple trajectories. Existing approaches to animal movement modeling mainly addressed the first objective and they are yet soon limited when used for simulation. Individual-based models based on ad-hoc formulation and empirical parametrization lack of generability, while state-space models and stochastic differential equations models, based on rigorous statistical inference, consist in 1st order Markovian models calibrated at the local scale which can lead to overly simplistic description of trajectories.
2. We introduce a ‘state-of-the-art’ tool from artificial intelligence - Generative Adversarial Networks (GAN) - for the simulation of animal trajectories. GAN consist in a pair of deep neural networks that aim at capturing the data distribution of some experimental dataset, and that enable the generation of new instances of data that share statistical similarity. In this study, we aim on one hand to identify relevant deep networks architecture for simulating central-place foraging trajectories and on the second hand to evaluate GAN benefits over classical methods such as state-switching Hidden Markov Models (HMM).
3. We demonstrate the outstanding ability of GAN to simulate ‘realistic’ seabirds foraging trajectories. In particular, we show that deep convolutional networks are more efficient than LSTM networks and that GAN-derived synthetic trajectories reproduce better the Fourier spectral density of observed trajectories than those simulated using HMM. Therefore, unlike HMM, GAN capture the variability of large-scale descriptive statistics such as foraging trips distance, duration and tortuosity.
4. GAN offer a relevant alternative to existing approaches to modeling animal movement since it is calibrated to reproduce multiple scales at the same time, thus freeing ecologists from the assumption of first-order markovianity. GAN also provide an ultra-flexible and robust framework that could further take environmental conditions, social interactions or even bio-energetics model into account and tackle a wide range of key challenges in movement ecology.

## 1 Introduction

Recent advances in telemetry and electronic systems enabled ecologists to track free-ranging animals and to gather large trajectories datasets (Ropert-Coudert *et al*., 2009; Bograd *et al*., 2010; Chung *et al*., 2021). GPS loggers have been at the forefront of this breakthrough, and can now provide precise and accurate data on the movements of many species, such as seabirds (Wakefield *et al*., 2009; Yoda, 2019). These movement data contain crucial information about animal behaviour including habitat selection, migration patterns, and foraging strategies but present key challenges for movement ecologists for elucidating underlying animal movement ecology (Hays *et al*., 2016).

Animal trajectories are generally seen as a succession of elementary behavioural events called steps (Nathan *et al*., 2008), and the use of random walk has received increased attention for describing the correlation and dynamics of such step sequences (Turchin, 1998; Codling *et al*., 2008). This includes correlated random walks (e.g. Bergman *et al*., 2000), Lévy random walks (e.g. Viswanathan *et al*., 2008), state-space models (e.g. Patterson *et al*., 2008) and stochastic differential equations models (e.g. Michelot *et al*., 2018), which offer different approaches to describing animal movement patterns. Random walks have also been used as “building blocks” for more complex models to simulate realistic global movement patterns of different animals. To this end, animal behavioural heterogeneity is often taken into account by developing state-switching models where animal movements are seen as the outcome of distinct behavioural states (e.g. travelling, resting and foraging) (Morales *et al*., 2004). This modelling approach enabled notably the simulation of central-place foraging trajectories (hereafter CPF), such as seabirds trajectories during breeding season (e.g. Boyd *et al*., 2016; Zhang *et al*., 2017) or marine mammals trajectories (e.g. Satterthwaite & Mangel, 2012; Massardier-Galatà *et al*., 2017).

Trajectory simulations models are indeed of crucial interest in the emerging field of movement ecology to assess the role of environmental heterogeneity, perceptual ranges, memory, and other mechanisms in creating different movement patterns (Avgar *et al*., 2013). It is also used to assess effectiveness of spatial management regimes and evaluate connectivity between different populations (Palmer *et al*., 2011; DeAngelis & Grimm, 2014). It may also serve as a null model for testing hypotheses concerning movement (Michelot *et al*., 2017). Yet, except few studies that fitted movement models for the simulation of CPF trajectories by likelihood maximization or through an approximate bayesian computation framework (Michelot *et al*., 2017; Zhang *et al*., 2017), most approaches consist in individual-based models relying on empirical parametrization and on ad-hoc mechanistic assumptions. There is therefore a growing interest for data-driven approach in ecology, and in particular for deep learning techniques (Malde *et al*., 2020), to better match the variability of available tagging datasets.

Deep learning refers to a neural network with multiple layers of processing units (LeCun *et al*., 2015). By decomposing the data into these multiple layers, deep neural networks allow to learn complex features for representing the data with high level of abstraction at multiple-scales. It has been widely used in ecology notably for segmentation, classification and identification tasks (Christin *et al*., 2019). Deep learning tools have yet recently demonstrated a great ability for simulating complex systems particularly using Generative Adversarial Networks (GAN) (Goodfellow *et al*., 2014). GAN consist in a pair of deep neural networks that aim at capturing the data distribution of some experimental dataset, and that enable the generation of new instances of data that share statistical similarity. It has recently become a state-of-the-art approach for generating various type of data such as image, audio, and spatio-temporal data including human trajectories (Cao *et al*., 2019; Gao *et al*., 2020).

This paper proposes therefore to introduce generative adversarial networks for the simulation of animal trajectories and in particular for the simulation of CPF trajectories. Our key contributions are the design of different GAN architectures, and the evaluation of GAN benefits over ‘state-of-the-art’ tools, i.e. state-switching Hidden Markov Models (HMM) over real seabird foraging trajectories. In particular, this study aim to demonstrate the ability of GAN to reproduce multiple scales at the same time, thus freeing ecologists from the assumption of first-order markovianity. We further discuss the potential use of GAN and the promise of such an approach for tackling nowadays ecological challenges.

## 2 Material and Methods

### 2.1 Generative Adversarial Network

#### 2.1.1 Background

The location of an animal is generally represented by a discrete-time or time-continuous stochastic process (*X_t_*)_*t*≥0_, where *t* denotes the time. A Markov hypothesis is classically stated for this movement process. It assumes that one can fully predict the distribution of the future state of an animal given its current state (Patterson *et al*., 2008). In a probabilistic framework, it consists in elucidating the density function *p*(*X*_*t*+1_|(*X_t_* = *x_t_*), *θ_t_*), where *θ_t_* depends on various parameters such as enviromental factors, motion capacity, navigation capacity or the internal state of the animal (Nathan *et al*., 2008). This Markovian hypothesis can only be regarded as an approximation of real movement patterns, which generally also involve long-term dependencies. The calibration of these models generally rely on the maximization of the likelihood of observed trajectories at local scales. It typically comes to estimating the joint distribution of step distance and heading turning angle from GPS tracks sampled at regular time intervals.

A wide range of probabilistic models can be restated as the composition of deterministic function *G* and of the sampler of a random latent variable. We may illustrate this point for correlated random walk (CRW) as presented in Patterson *et al*. (2008). Let is denote by *s_t_* and *ϕ_t_* the step length and the heading at time *t*. The CRW can be written as:

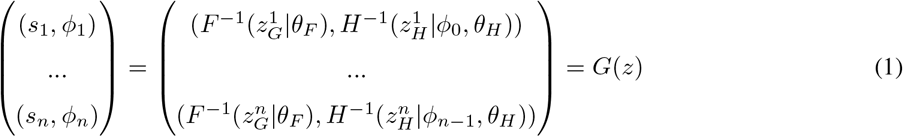

where *F* and *H* are cumulative density functions with parameters *θ_F_* and *θ_H_*, often chosen as Log-Normal and Von Mises distributions, and where 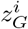 and 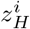 are independent samples from the uniform distribution over [0, 1].

The generative model in a GAN also relies on the application of a deterministic function *G* to random samples of a latent variable *z* according to a predefined distribution. Function *G* is chosen within a parametric family of differentiable functions and implemented as a neural network for flexibility. The other major difference with statistical inference approaches classically exploited in movement ecology lies in the calibration approach from data. Rather than stating the calibration as the maximization of a likelihood criterion, the calibration of the generator of a GAN involves the simultaneous training of another deep network *D* (referred as the discriminator) that learns how to distinguish simulated data (i.e. *G*(*z*)) from real data. The architecture of GAN is given in Fig. 1A. If no disriminator can distinguish the simulated and real data, it means that the generator truly sample the unknown distribution of the training dataset (Goodfellow *et al*., 2014).

**Figure 1:**
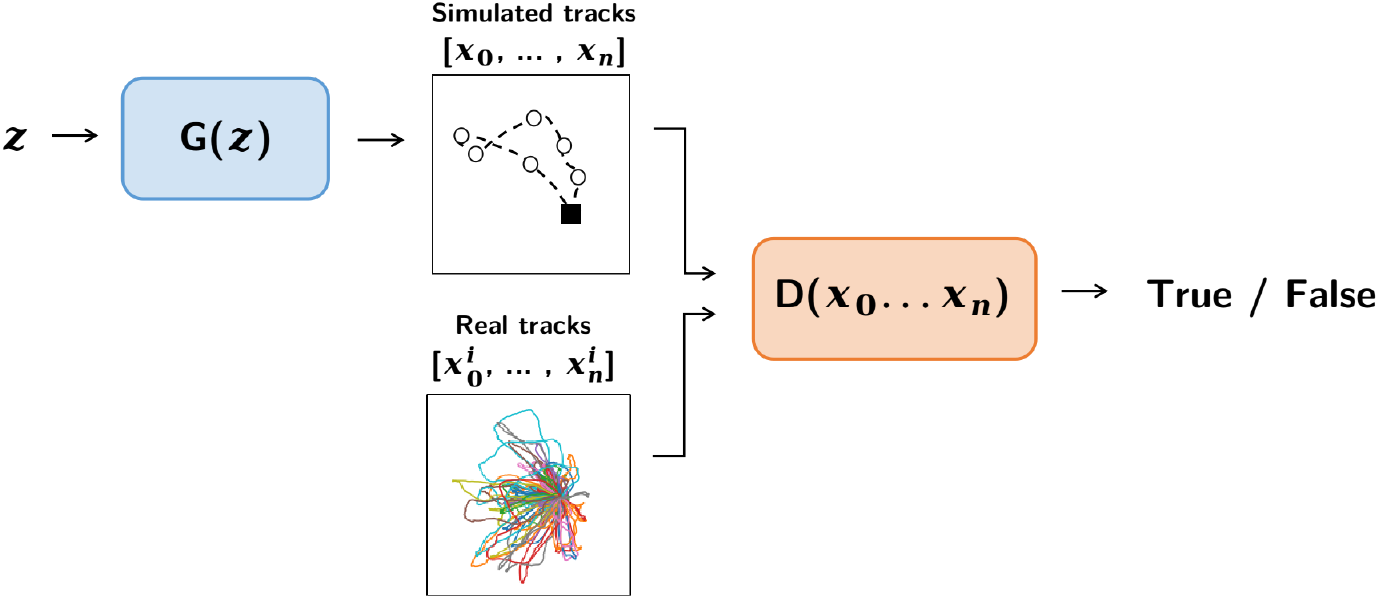
GAN Architecture: Global architecture of a generative adversarial network. *G* refers to the generator network that takes as input a random noise vector *z* and outputs a trajectory *x*. *D* is the discriminator network that aims to distinguish real trajectories from simulated ones

#### 2.1.2 Network Architecture

Numerous deep networks architecture can be used for both generator and discriminator networks. Long short-term memory (LSTM) networks and convolutional neural networks (CNNs) are probably the most popular, efficient and widely used deep learning techniques (Alom *et al*., 2019). In this study we used two generator and two discriminator architectures, in both cases one architecture relies on CNN while the second one is based on LSTM (see Fig. 2). Here, we briefly present the motivation and functioning of these networks. We refer the reader to Christin *et al*. (2019) for a detailed introduction to deep networks.

**Figure 2:**
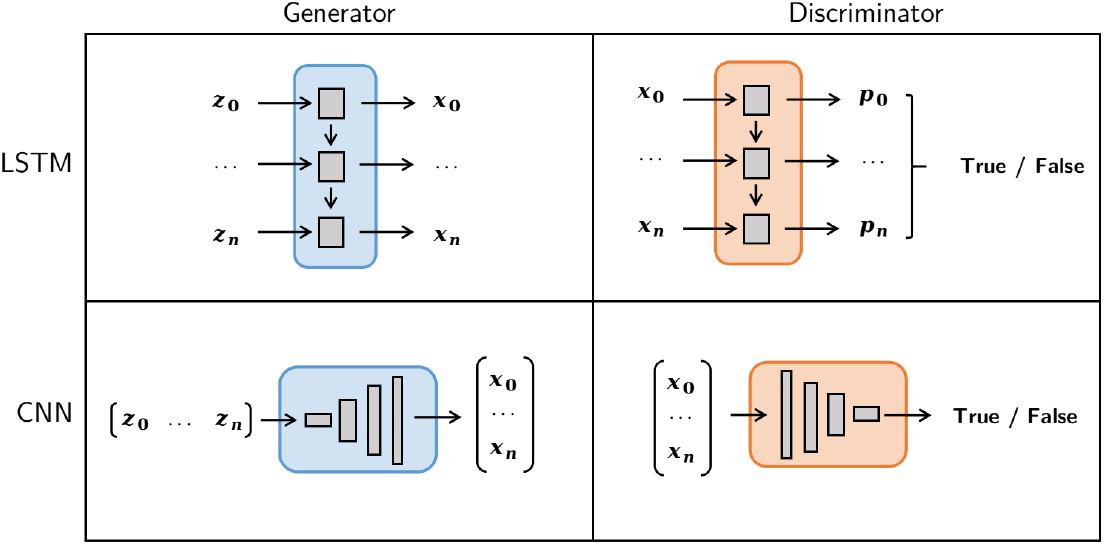
LSTM vs CNN. Architecture of LSTM and CNN networks used in this study.

##### LSTM

Long Short Term Memory (LSTM) Networks are among the state-of-the-art architectures of recurrent neural networks dedicated to the modeling and processing of time series. LSTM then seems a natural framework for the simulation of trajectories. A key feature of LSTM is its ability to identify and exploit long-term dependencies through gating processes (Hochreiter & Schmidhuber, 1997). Various studies that explored deep learning for the analysis of animal trajectories relied on LSTMs (Wijeyakulasuriya *et al*., 2020; Li *et al*., 2021). LSTM-based architecture have also been used in numerous recurrent GAN for music, and medical time-series generation (Mogren, 2016; Esteban *et al*., 2017).

In our study, we used a generator network composed of a LSTM layer that takes a different random seed at each temporal input, and produces a sequence of hidden vectors with 16 features. These hidden vectors encode the state of the trajectory. An additional dense layer maps the 16-dimensional hidden vector at a given time step to the corresponding longitudinal and latitudinal displacements. The cumulative sum of these elementary displacements form the last layer of generator to put a time series of positions (see Fig. 2).

We can also exploit a LSTM for the discriminator. Given a sequence of positions (longitude, latitude), the LSTM acts as an encoder of this sequence in some higher-dimensional latent space. A dense layer was then applied to assign a probability of being realistic at each position of the sequence. Overall, the output of the discriminator is the associated mean probability to assess the quality of the whole trajectory (see Fig. 2).

##### CNN

CNN architectures exploit convolutional layers and are the state-of-art architectures for a wide range of applications, especially for signal and image processing tasks (Alom *et al*., 2019). They are particularly effective at extracting low-level and high-level features from n-dimensional tensors, and have reecntly been illustrated for animal trajectory segmentation (Roy *et al*., 2021).

CNN are also widely exploited in GANs (Radford *et al*., 2016), and have been eventually used for spatio-temporal data generation (Gao *et al*., 2020). Here, we follow the general architecture proposed in Radford *et al*. (2016) for image generation. The generator takes as input a random noise vector that can be seen as a latent representation of a global time-series. It then applies a series of successive fractional-strided convolutions to map the latent representation into time-series with increasing number of points and decreasing features, until it outputs a 2-dimensional vector of the required length (see Fig. 2). In our work, we used a batchnorm and a ReLU activation after every fractional-strided convolutions, except for the output that used only a Tanh, as suggested by (Radford *et al*., 2016). We may point out that, in this CNN architecture, there is explicit sequential modeling of the trajectory and the latent representation may not be a time-related.

Regarding the CNN-based discriminator, we also applied successive strided convolutions in order to transform the initial trajectory into time-series with decreasing lengths and increasing numbers of features, until we obtained a latent vector describing the whole trajectory. We used batchnorm and LeakyReLU activation after every strided convolutions as suggested by (Radford *et al*., 2016). The last layer is a dense layer with a sigmoid activation that transforms the latent representation into a probability for the trajectory of being realistic (see Fig. 2).

#### 2.1.3 Adversarial training and spectral regularization

For a given architecture, networks’ parameters are estimated using adversarial training, i.e. the two networks compete in a minimax two-player game given by Eq. 2. Discriminator *D* is trained to maximize the probability of assigning the correct label to both training examples and samples from *G*, i.e. to maximize log *D*(*x*) + log(1 – *D*(*G*(*z*))). Generator *G* is simultaneously trained to fool the discriminator, i.e. to minimize log(1 – *D*(*G*(*z*))).

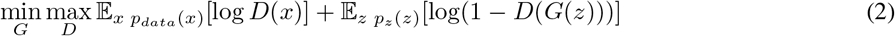

Numerically, we apply stochastic gradient descents over the discriminator and generator successively where at each iteration, we compute the training losses for a randomly sampled subset of *m* trajectories within the training dataset^1^:

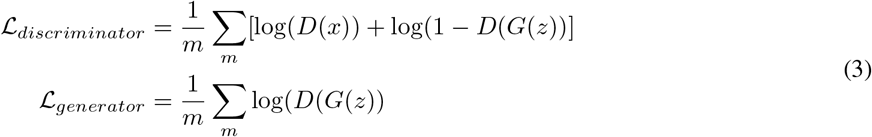

We may complement the training loss of the generator with additional terms, including both application-specific (Ledig *et al*., 2017) and regularization (Durall *et al*., 2020) terms. In particular, recent studies have demonstrated that a spectral regularization may have positive effects both on the training stability and output quality of generative networks for image simulation (Durall *et al*., 2020). We tested here a similar approach with the following spectral loss *L_spectral_* to the generator’s gradient descent:

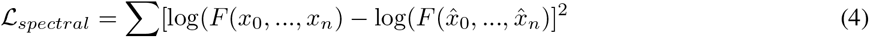

Where *F* is the module of the Fourier Transform of a 2-dimensional time-series, *x* and 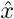 are real and simulated trajectories respectively.

### 2.2 Case studies and experiments

#### 2.2.1 Datasets

GPS were fitted to tropical boobies during breeding period from two distinct locations (Table 1). Trajectories consist in foraging trips where seabirds look for preys at sea and come back to their colony. Data points have been linearly re-interpolated at regular time steps, and coordinates have been centered on the colony’s location and reduced. Finally, trajectories have been padded with zeros so that all longitude/latitude time-series from a dataset would have the same length. Example of real trajectories are given in Fig. 6.

**Table 1:** Datasets Overview

| Species | Country | Nb of trips | Resolution | Padding |
| --- | --- | --- | --- | --- |
| *Sula sula* | Brazil | 30 | 1h | 20 steps |
| *Sula dactylatra* | Brazil | 50 | 15s | 200 steps |
| *Sula variegata* | Peru | 78 | 5s | 200 steps |

#### 2.2.2 Architecture selection experiment

We first designed an experiment to compare different GAN architectures. For this experiment, we considered the simplest dataset with a 1 hour time resolution (see Table 1). All trajectories involved 20 time steps. We evaluated four different GANs corresponding to every generator-discriminator pairs for the considered CNN and LSTM architectures: e.g., we call ‘LSTM-CNN’ the GAN with a LSTM network as generator and a CNN as discriminator.

For all generators, the input random noise vector consisted in 20 samples from a uniform distribution on [0,1]. All networks had about 1500 parameters, and details on network structure are available on our github repository^2^. We trained all networks over 5000 epochs with a learning rate of 2e-4 using the loss functions given in Eq. 3. The score of each approach was assessed by computing the mean squared error of the logarithmic Fourier decomposition spectrum of simulated and real trajectories, 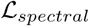 (see Eq. 4).

#### 2.2.3 GAN vs HMM experiment

In this section we compared the best GAN architecture from the previous experiment, namely ‘CNN-CNN’ GAN architecture, to the state-of-the-art approach to animal trajectories simulation, i.e. state-switching Hidden Markov Models (HMM). We tested both methods on the two datasets with 200-step time-series consisting in trajectories of tropical boobies from two completely distinct ecosystems and with different foraging strategies (see Table 1).

##### GAN

The input random noise vector consisted in 256 samples from a uniform distribution on [0,1]. We trained the ‘CNN-CNN’ GAN architecture separately on each dataset over 5000 epochs and with a learning rate of 2e-4. We used a spectral regularization to better reproduce the spectral features of real trajectories, especially for fine time scales, and to increase learning stability. Details on the structure is available on our github repository^1^.

##### HMM

For comparison we fitted a ‘state-of-the-art’ state-switching HMM to seabirds CPF trajectories. We followed the methodology presented by (Michelot *et al*., 2017), which relies on a rigorous statistical inference.

Movements were described as a sequence of step lengths and turning angles that we fitted with gamma distribution and von Mises distribution respectively. Three behavioural states were used for the Peruvian datasets i.e., “searching”, “foraging” and “inbound”, while a fourth state was added with the Brazilian dataset i.e., “resting” (Fig. 6). For states “searching”, “foraging” and “resting”, we described movement as correlated random walks (CRW), while for state “inbound” we used a biased random walk (BRW) with attraction toward the colony. In order to force the return to the colony, we fixed some terms of the transition matrix thus ensuring that the sequence of states alternates first with “searching”, “foraging” and “resting”, and is then forced to stay in state “inbound”.

These state-switching HMM were fitted to real data according to a maximum likelihood criterion. Fitted models were used to simulate trajectories. The initial step was sample from real data, and we iteratively sampled next steps, until the trajectory went back to the colony. In practice, we stopped the simulation once a location was simulated within a 1-km radius around the colony.

##### Implementation details

GAN were implemented and trained using Pytorch (Paskze *et al*., 2019). HMM were fitted using the momentuHMM R package (McClintock & Michelot, 2018). The code of all the reported experiments is available on our github repository: https://github.com/AmedeeRoy/BirdGAN

## 3 Results

### 3.1 Architecture selection experiment

From the four GAN architectures the fully deep convolutional GAN lead to the best results with better convergence and lowest computation time (Fig. 4 and Table 2). GANs with LSTM-based discriminators seemed particularly unstable with highly variable performance through epochs (4). Importantly, only GANs with CNN-based generators managed to simulate looping trajectories. For instance, the ‘LSTM-CNN’ GAN generated relatively good trajectories with a spectral error 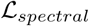 lower than 3, yet without being able to loop (Fig. 3).

**Figure 3:**
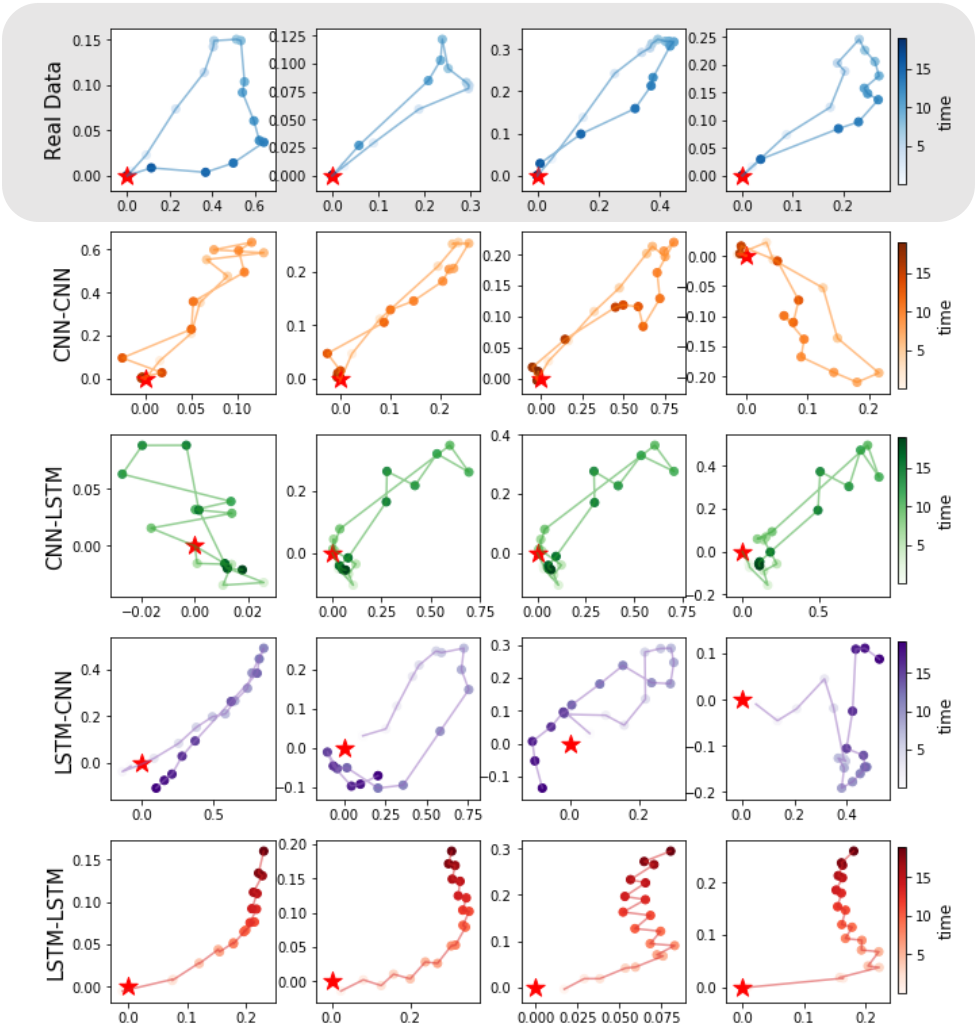
Real vs Simulated Trajectories: 4 examples of trajectories generated by each GAN architecture tested on the 20-step dataset (see Table 1). The four different GANs correspond to every generator-discriminator pairs for the considered LSTM and CNN architectures: e.g., we call ‘LSTM-CNN’ the GAN which has a LSTM Network as generator and a CNN as discriminator.

**Figure 4:**
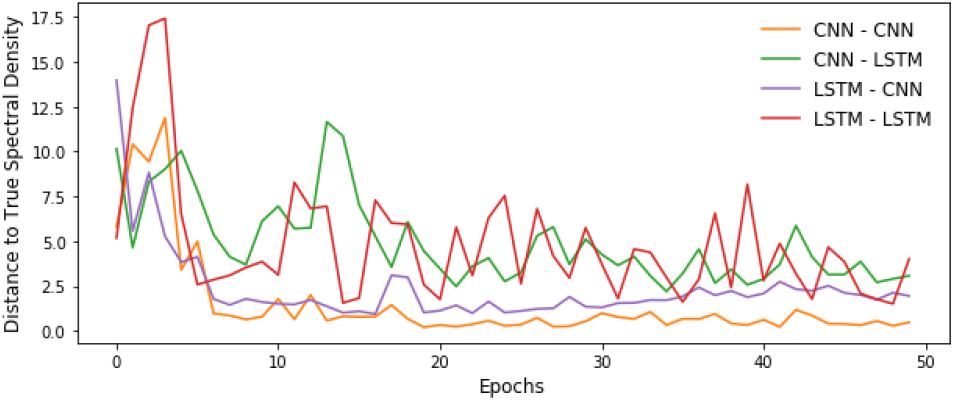
Convergence of GAN architecture over 5000 epochs: The four different GANs correspond to every generator-discriminator pairs: e.g., we call ‘LSTM-CNN’ the GAN which has a LSTM Network as generator and a CNN as discriminator. Distance to true spectral density is computed with Eq. 4

**Table 2:**
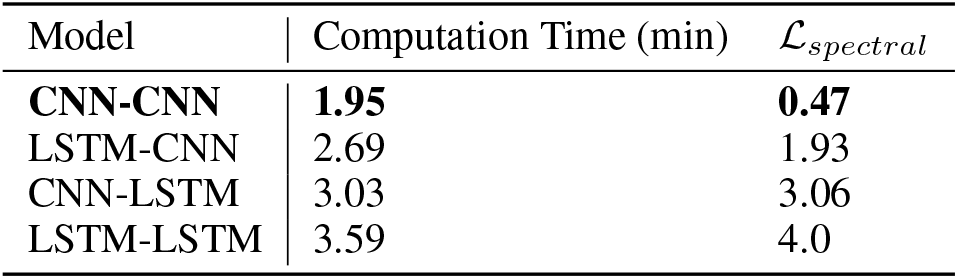
Comparison of GAN architectures

### 3.2 GAN vs HMM experiment

On both datasets, GAN and HMM managed to converge and to simulate relatively ‘realistic’ CPF trajectories (see simulated trajectories in Fig. 6). However, the spectral distribution of GAN-derived synthetic trajectories matched better the spectral distribution of real trajectories (Fig. 5). In particular, the mean spectral error 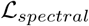 was about 4 times smaller using GAN than using HMM (Table 3). This was particularly highlighted for the highest frequencies (Fig. 5). On the Peruvian dataset, HMM failed to reproduce spectral distributions both at lower and higher frequencies (Fig. 5A), and on the Brazilian dataset, it failed in the higher frequencies only (Fig. 5B). By contrast, HMMs outperformed GANs to sample relevant step distributions (Fig. 7).

**Figure 5:**
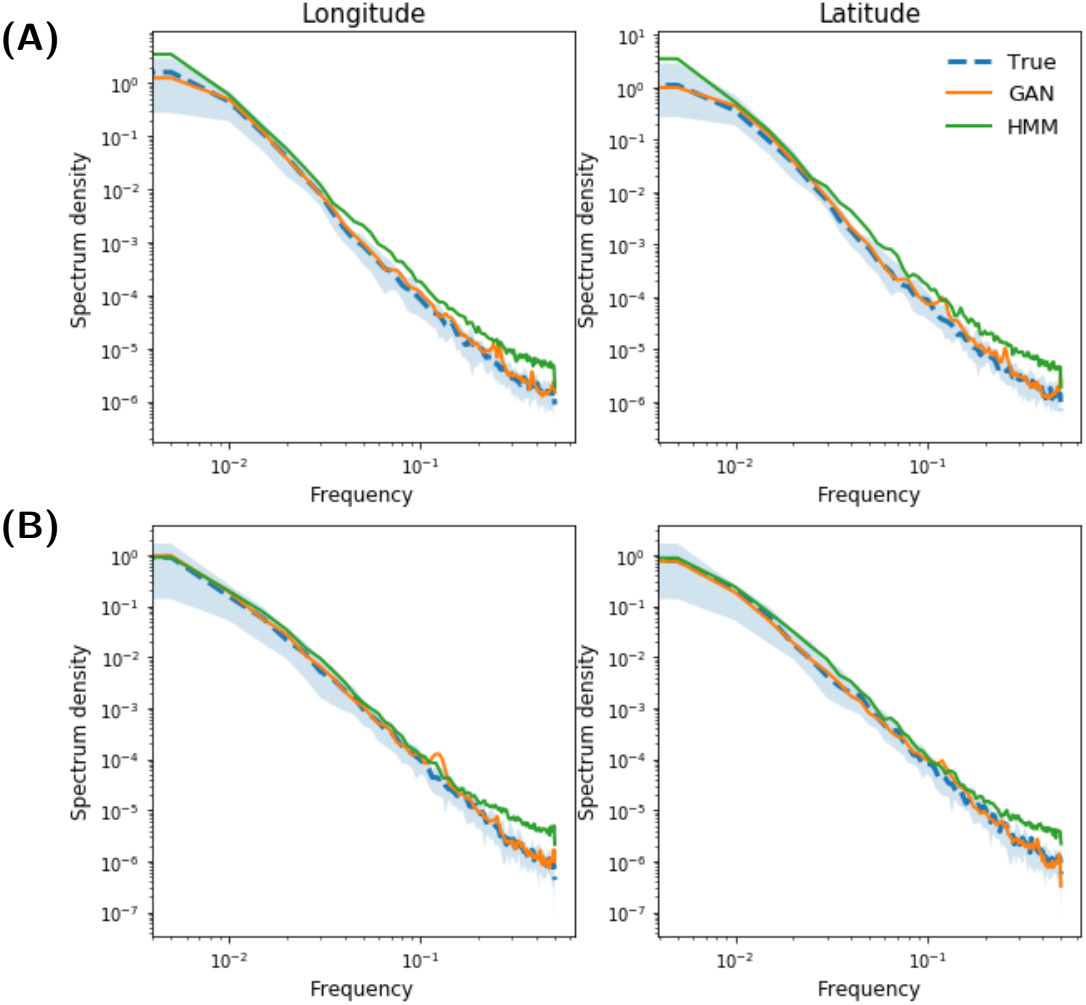
Mean Fourier Spectrum of real trajectories used for training (blue), synthetic trajectories generated by a ‘CNN-CNN’ GAN (orange) and synthetic trajectories generated by HMM (green). (A) is for the 200-step Peruvian dataset, and (B) is for the 200-step Brazilian dataset (see Table 1). In these plots, we computed the mean Fourier spectrum for datasets of 100 simulated trajectories.

**Figure 6:**
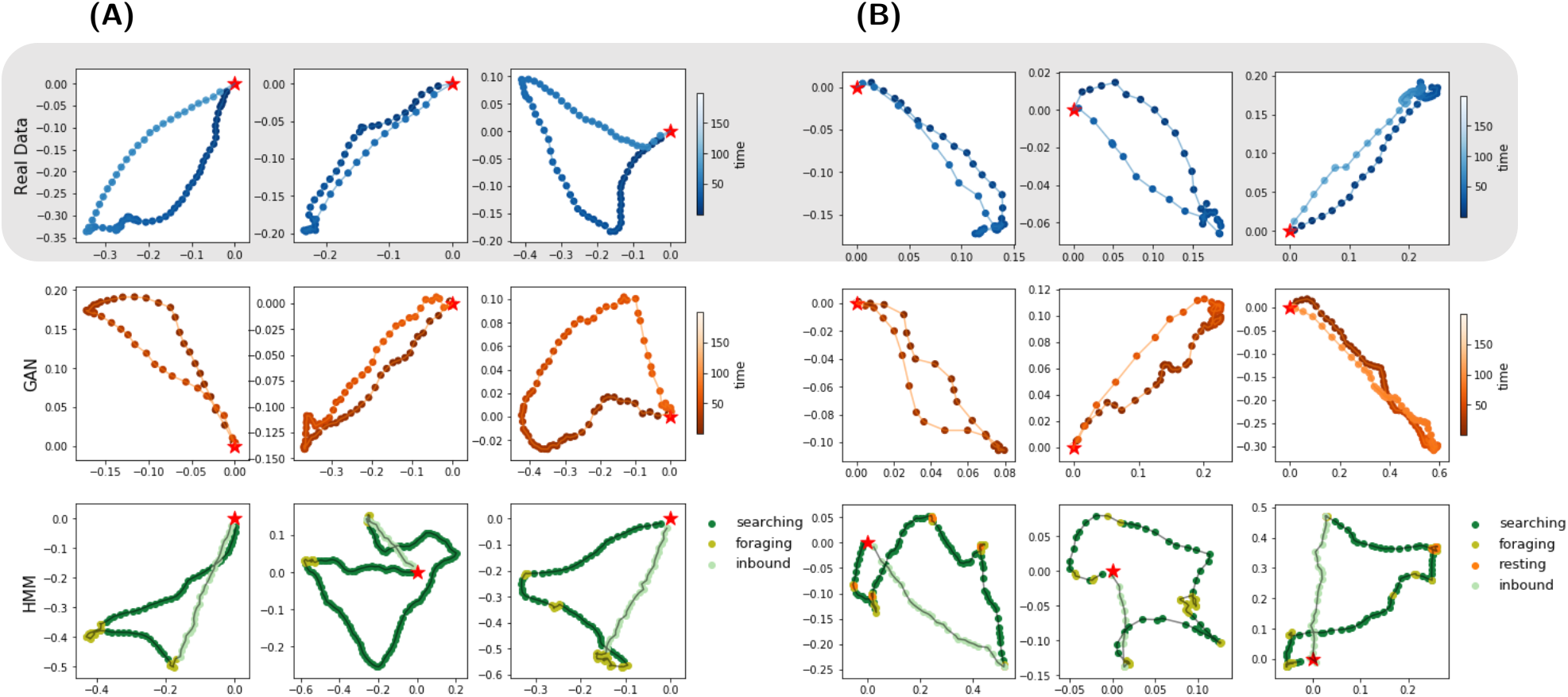
Example of trajectories. Real trajectories used for training are in blue, synthetic trajectories generated by a ‘CNN-CNN’ GAN are in orange and trajectories generated by HMM are in green. (A) is for the 200-steps Peruvian dataset, and (B) is for the 200-steps Brazilian dataset (see Table 1)

**Figure 7:**
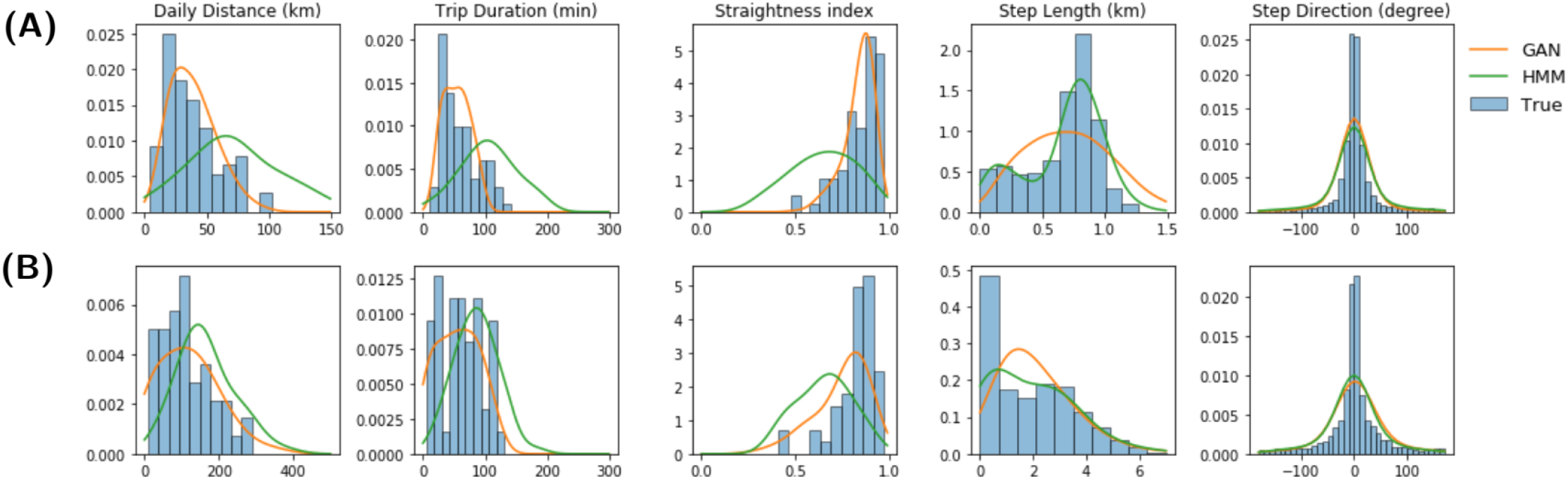
Histogram of descriptive statistics. derived from real trajectories used for training (blue), 100 synthetic trajectories generated by a ‘CNN-CNN’ GAN (orange) and 100 synthetic trajectories generated by HMM (green). (A) is for the 200-steps Peruvian dataset, and (B) is for the 200-steps Brazilian dataset (see Table 1)

**Table 3:**
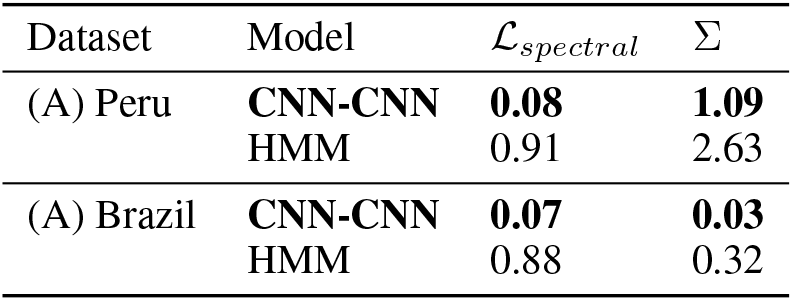
Properties of GAN and HMM simulations. 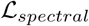 stands for the mean squared error of the logarithmic Fourier decomposition spectrum presented Fig. 5 and Σ stands for the mean squared error of the position distributions presented Fig. 8

| Dataset | Model | $\mathcal{L}_{spectral}$ | $\Sigma$ |
| --- | --- | --- | --- |
| (A) Peru | **CNN-CNN** | **0.08** | **1.09** |
|  | HMM | 0.91 | 2.63 |
| (A) Brazil | **CNN-CNN** | **0.07** | **0.03** |
|  | HMM | 0.88 | 0.32 |

Yet, GAN models better capture the real data distribution as it is able to simulate a set a trajectories that has similar global statistics the reference dataset does. For instance, our synthetic trajectories have consistent trip distance, trip duration and the straightness index distributions (see Fig. 7). The straightness index of a trajectory is defined as two times the quotient between the max range to the colony and the trip total distance and is a proxy for tortuosity (Benhamou, 2004). The trained GANs also capture spatial information as they reproduce position distributions of observed trajectories (Fig. 8 and Table 3). GAN-derived synthetic trajectories were indeed mainly heading toward some area of interest (i.e. westward to the colony on the Peruvian dataset, and to the north-east and south-east of the colony on the Brazilian dataset), while HMM-derived trajectories are uniformly directed to all directions around the colony.

**Figure 8:**
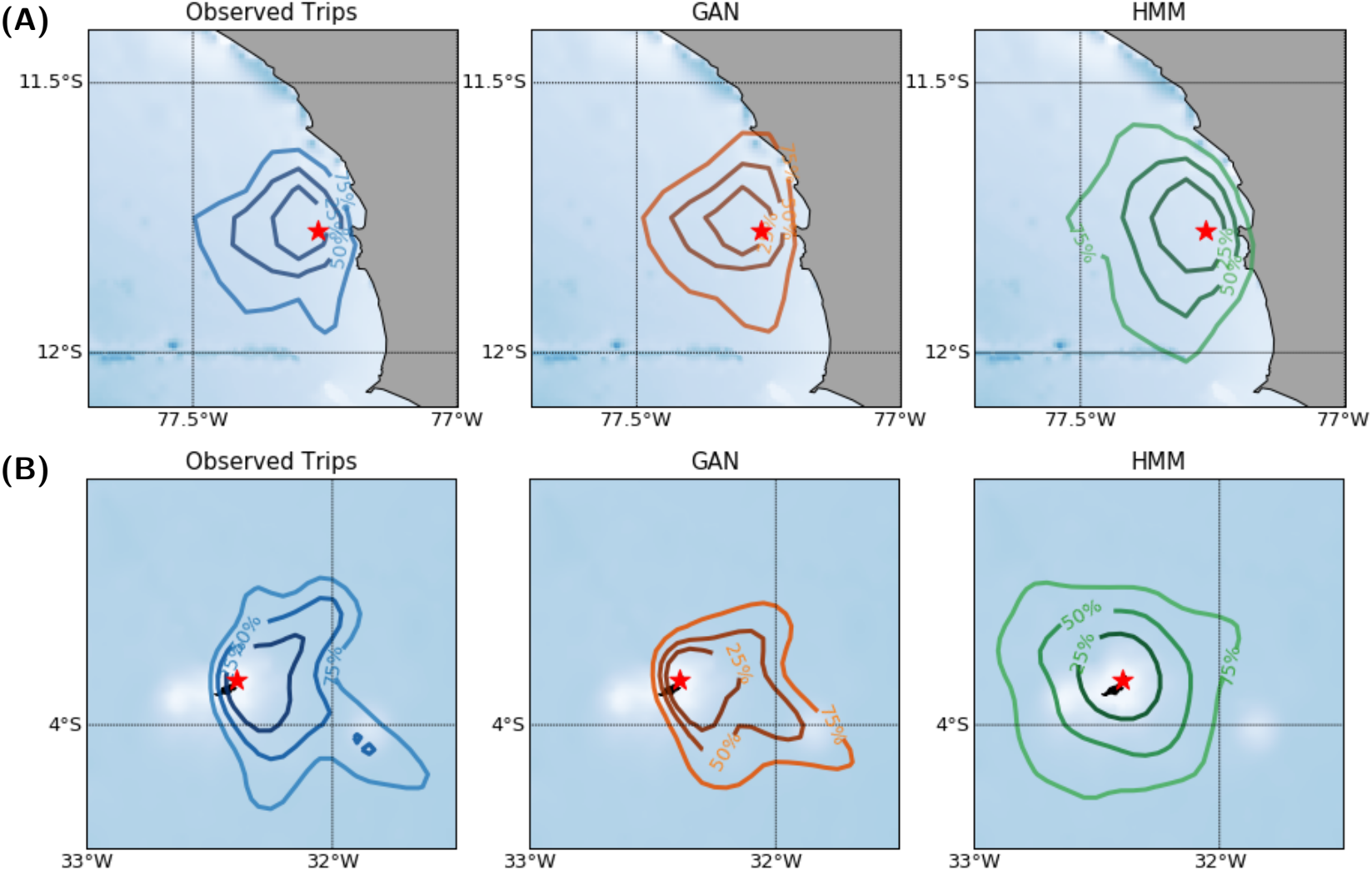
Position Distribution. Kernel Density Estimation of position distributions on real trajectories used for training (blue), 100 synthetic trajectories generated by a ‘CNN-CNN’ GAN (orange) and 100 synthetic trajectories generated by HMM (green). (A) is for the 200-steps Peruvian dataset, and (B) is for the 200-steps Brazilian dataset (see Table 1)

## 4 Discussion

Deep learning has become the state-of-the-art framework for a wide range of problems in ecology such as classification and segmentation tasks mainly for image analysis (Weinstein, 2018; Christin *et al*., 2019). Applications to trajectory data (Browning *et al*., 2018; Peng *et al*., 2019; Roy *et al*., 2021) have also recently emerged. Despite recent advances in deep learning for the simulation of complex systems, few studies have explored generative models, and particularly of Generative Adversarial Networks (GAN) or Variational Auto-Encoders (VAE) for simulating ecological data. To our knowledge, deep convolutional GAN have been used only for data augmentation in simulating plant or insect images so far (Giuffrida *et al*., 2017; Lu *et al*., 2019; Madsen *et al*., 2019; Silva *et al*., 2021). Our study demonstrates that GANs are also great tools for the generation of other ecological data such as animal trajectories.

GANs showed their great ability to capture the trajectory data distribution, except for first-order step distribution. In opposition, the current state-of-the-art approaches such as multi-state HMM are calibrated at a local scale and are unable to bring out global patterns from these local features. Our numerical experiments pointed out that the relationship between local and global features may be complex for real trajectory data. GANs are explicitly trained so that they best reproduce the characteristic multi-scale features of real trajectories. Through strided convolutions, the considered CNN discriminator likely overlooks the highest frequencies to focus on larger-scale information. Besides, the CNN generator does not explicitly represent a trajectory as a sequential process, which may also impede its ability to reproduce well step distribution. This may be a general property of convolution GAN architectures. For instance, GANs for image generation including object appearance but may not simulate realistically fine-scale textures Cao *et al*. (2019). Future work could further investigate new GAN architectures to address this issue. The combination of CNN architectures to sequential ones such as LSTM-based architectures appears as a natural research direction to explore.

We believe our study will open new research avenues for the exploration of ecological questions using GANs through both generator and discriminator networks. The generator network is a sampler of trajectory data, and it could be used as a null generative model for testing ecological hypothesis such as segregation of foraging areas (Bolton *et al*., 2019), individual foraging site fidelity (Owen *et al*., 2019), or for generating relevant pseudo-absence in order to calibrate some ecological niche model (Hückstädt *et al*., 2020). By computing the probability of being a ‘realistic’ trajectory, the discriminator network provides a metric of data similarity, and it could be used within comparative study of foraging strategies in order to assess sex-specific (Lewis *et al*., 2005), breeding stage (Lerma *et al*., 2020) or inter-colony differences (Harding *et al*., 2013).

Numerous existing varieties of GAN could also provide a great support for movement ecology, such as conditional GAN (Isola *et al*., 2018). A conditional GAN consists in a GAN with some external variable that is supposed to condition GAN’s output. It could therefore be possible to test for condition that would explain behavioural variability such as individual characteristics (e.g. sex, mass, breeding stage), or environmental characteristics (e.g. prey distributions, oceanographic features). Testing different environmental scenarios and predicting associated animal trajectories is indeed a topic of interest notably for predicting the potential impact of climate change on animal behavioural (Hückstädt *et al*., 2020). Conditional GANs could also be applied to the interpolation of trajectory data and to produce superresolution trajectories as performed in computer vision (Ledig *et al*., 2017).

Increasing literature concerns also the use of physics-informed GANs for modeling dynamic systems that aims at encoding known physical laws into the framework of GANs. This can be achieved either by encoding directly stochastic differential equations into the architecture of generators (Yang *et al*., 2018) either in incorporating additional penalty terms into the optimization loss function of GANs (Bode *et al*., 2019; Wu *et al*., 2020). Such approach might therefore enable ecologists to link GAN-type approaches to a mechanistic formulation of animal movement for instance by including bio-energetics equations, animal perceptions or cognitive relationships in their movement processes in a straightforward manner.

GAN provide an ultra-flexible framework where traditional methods such as HMM struggle at accounting for environmental heterogeneity, perceptual ranges, memory, and social interactions and can suffer from computational time when maximizing likelihood of a complex state-space model. This study introduces therefore a truly promising tool that would allow to simulate free-ranging animal movement and freeing movement ecologists from 1st order Markov property that often lead to overly simplistic description of animal behaviour.

## Acknowledgment

This work is a contribution to the TRIATLAS project (European Union’s Horizon 2020 research and innovation program – grant agreement No. 817578), to the Mixed International Laboratory TAPIOCA Program, and to the Young Team IRD Program (JEAI) TABASCO. RF was supported by LEFE program (LEFE MANU project IA-OAC), CNES (grant SWOT-DIEGO) and ANR Projects Melody and OceaniX. Fieldworks have been conducted thanks to the cooperative agreement between IRD, the Agence Nationale de la Recherche (ANR) project TOPINEME, and of the International Joint Laboratory DISCOH. We would like thank all people involved in fieldworks including french, Brazilian and Peruvian researchers/students. We thank the Brazilian Ministry of Environment and Fernando de Noronha’s firemen for the authorization and technical support to capture seabirds in Brazil. We thank the Ministry of Agriculture of Peru and island guards for the authorization and technical support to capture seabirds in Peru.

## Authors’ contributions

All authors conceived the ideas. A.R. developed the methodology and wrote R and python codes. All authors contributed critically to the writing of the article. Authors declare that there is no conflict of interests.

## Data avaibility

All data and codes are available on our github repository https://github.com/AmedeeRoy/BirdGAN.

## Footnotes

1 This subset is referred to as a batch in the deep learning literature.

2 https://github.com/AmedeeRoy/BirdGAN

## Notes

https://github.com/AmedeeRoy/BirdGAN

### Competing Interest Statement

The authors have declared no competing interest.

